# Neurexin mediates neuropeptide release from cholinergic motor neurons through dense-core vesicle localization

**DOI:** 10.64898/2026.08.31.748337

**Authors:** Vina Tikiyani, Michael P. Hart

## Abstract

Neurexins are critical synaptic cell adhesion molecules that play many roles in modulating neurotransmitter release and synaptic function, and are high-confidence risk genes for neurodevelopmental conditions such as autism. Understanding the function of the neurexin superfamily has been challenging in mammalian systems that have 3 genes (*NRXN1-3*) that encode 2-3 major isoforms, which undergo extensive alternative splicing and generate thousands of transcripts. In contrast to mammals, *C. elegans* has a single gene, *nrx-1*, encoding only long α and short γ isoforms. Neurexins canonically regulate synapse morphology and function in a neuron- and context-specific manner, through mechanisms related to release of chemical neurotransmitters and receptors. Whether neurexins (*nrx-1*) impact other secretory molecules such as neuropeptides (NPs) and NP containing dense-core vesicles (DCVs) is not well understood. Here, we report that loss of *nrx-1* increases the release of multiple NPs from cholinergic motor neurons in *C. elegans*. Using tissue specific expression and degradation of endogenous NRX-1, we find that *nrx-1* functions in NP release from cholinergic neurons in a cell-autonomous manner. We confirm that loss of *nrx-1* impacts cholinergic active-zone number, but also find it regulates the clustering, distribution, and expression of the DCV protein, IDA-1 (PTPRN), and the DCV secretion regulator, UNC-31 (CADPS). We find that *nrx-1* functions to maintain separation and juxtaposition of neurotransmitter and NP release sites and DCV localization. Loss of cholinergic excitation (*unc-17*) or GABAergic inhibition (*unc-25*) did not impact cholinergic NP release, but that the increased NP release upon loss of *nrx-1* is dependent on the calcium channel *unc-2*. We find that neurexins can regulate NP signaling, a novel mechanism to modify circuits and behaviors, and of potential importance for *NRXN1* associated human conditions.

## INTRODUCTION

Neurons communicate through multiple modalities that differ in timescale, distance, and downstream responses. These include direct coupling through gap junctions, fast neurotransmission at the synaptic cleft (GABA, glutamate, acetylcholine), and slower, further-acting neuromodulation by biogenic amines and neuropeptides. These signaling modalities act together to tune neuronal and circuit properties across time and context to generate and modify behaviors. Neurotransmitters and neuropeptides are both released by SNARE-dependent, calcium-triggered fusion of vesicles. However, clear-core synaptic vesicles (SVs) are recycled locally and fuse at the active zone within milliseconds, whereas dense core vesicles (DCVs) are generated at the *trans*-Golgi network, cannot be refilled locally, and fuse at sites that remain poorly defined. How, when, and where DCV release is controlled, and how much of that control is shared with SVs, is not well understood.

Synaptic adhesion molecules are central regulators of neuronal communication and have functions regulating gap junctions and SV release at active zones. Neurexins are the canonical example: presynaptic cell adhesion molecules that organize the active zone, couple calcium channels to release sites, and set release probability through interactions with many synaptic ligands (Hu et al. 2012; Padmanabhan et al. 2026). Across systems, however, neurexin function has been defined almost exclusively through clear-core SVs with comparatively limited evidence for roles in monoamine or DCV pathways. In mouse serotonergic (5-HT) neurons, selective deletion of *Nrxn1–3* reduced evoked 5-HT release, serotonergic neuron numbers, and serotonin transporter expression (Cheung et al. 2023). Evidence linking neurexins to the DCV pathway is also emerging; deletion of β-neurexins in mice reduced DCV abundance in hippocampal and cerebellar presynaptic boutons without detectable alterations in Golgi ultrastructure or DCV-associated markers (Ferdos et al. 2022), However, whether this reduction in DCV abundance translates into altered neuropeptide release was not directly tested and whether neurexins directly regulate neuropeptide secretion remains unresolved. The known regulators of DCV release are primarily cytoplasmic (e.g. CADPS/UNC-31, MUNC13, RIM, RAB2/RAB10, synapsin), and the roles of trans-synaptic and adhesion molecules in neuropeptide release are less well known.

The three human neurexin genes each produce multiple isoforms from alternative promoters and start sites that, together with extensive alternative splicing, generate hundreds to thousands of distinct neurexin proteins (Rowen et al. 2002). This diversity likely underlies the range of neurexin function across neurons, circuits, and behaviors, and may also contribute to the phenotypic breadth of *NRXN1* loss in humans that can include autism, schizophrenia, Tourette syndrome, and seizure disorders. This breadth in behavioral phenotypes is difficult to account for through changes in fast synaptic transmission alone, and one in which neuropeptide systems are independently implicated. Defining the full set of mechanisms by which neurexins regulate neuronal communication, including neuromodulatory signaling, is therefore necessary to connect neurexin genotype to circuit and behavioral phenotype.

*C. elegans* carries a single neurexin gene, *nrx-1*, which regulates active zone and synapse development and function, and which has served as a discovery model for conserved neurexin mechanisms. Mechanistic findings on the γ neurexin isoform in synaptic development and function in *C. elegans* were recently recapitulated in mammals (Kurshan et al. 2018; Padmanabhan et al. 2026). We have reported roles for *nrx-1* isoforms in multiple *C. elegans* behaviors, including social feeding (Cowen et al. 2024), in which animals aggregate with one another, often near the edge of the bacterial lawn (“bordering”), which is characteristic of strains carrying the ancestral/social *npr-1* variant (de Bono and Bargmann 1998). Social feeding depends on gap junction and neuropeptide signaling, and we found an additional requirement for neurexin and classical neurotransmission, raising the possibility that neurexin also acts on neuropeptide signaling itself. *C. elegans* ventral nerve cord cholinergic motor neurons offer a tractable system in which to test this: they release both acetylcholine and NLP-class neuropeptides, they signal to both muscle and GABAergic motor neurons, their DCVs can be visualized and quantified *in vivo* with endogenous markers, and neuropeptide release can be measured quantitatively using a coelomocyte uptake of secreted fluorescent neuropeptide reporters.

Here we report that neurexin functions to restrict neuropeptide release. Loss of both NRX-1 isoforms increases release of multiple NLP-class neuropeptides from cholinergic motor neurons (*nlp-21*, *nlp-9*), and the conserved CCK-like *nlp-12*, while loss of just the longer α isoform is dispensable for restricting *nlp-21* release. Loss of *nrx-1* also increases the levels, but not number, of NLP-21 puncta in the axons of these same neurons. Targeted degradation of endogenous NRX-1 localizes this function to neurons, and expression of either NRX-1 isoform pan-neuronally or in cholinergic motor neurons restores normal release, indicating a cell-autonomous mechanism. Endogenously tagged NRX-1 shows colocalization with cholinergic motor neuron DCVs, which are organized into puncta juxtaposed to synaptic active zones (marked by RIMS/UNC-10). Loss of *nrx-1* alters active zone morphology, as previously reported, but also reduces DCV content and puncta number. This phenotype is not a secondary consequence of altered excitatory or inhibitory transmission, as neither *unc-17*/vAChT nor *unc-25*/GAD mutants reproduce it. Instead, NLP-21 release tracks with presynaptic calcium availability, as loss of *unc-2*, encoding a voltage-gated calcium channel, significantly reduces NLP-21 release and UNC-2/CaV2 and NRX-1 function together to regulate calcium-dependent NLP-21 secretion. Together, these results identify neurexin as a cell-autonomous regulator of neuropeptide release and signaling, and extend its function beyond fast synaptic transmission to another major neuronal signaling modality. This has important implications for how DCV and neuropeptides are regulated, for the role of synaptic adhesion molecules in this regulation, and for the conditions and disorders associated with disruption in neurexins and related genes.

## RESULTS

### *nrx-1* regulates the secretion of multiple neuropeptides from cholinergic motor neurons

Neurexins regulate the release of classical chemical neurotransmitters in clear core vesicles through many mechanisms, but whether they regulate the release or expression of other secretory molecules or dense core vesicles is not known. One recent study indicates that *Nrxn1* could regulate the number of neuropeptide containing dense-core vesicles (Ferdos et al. 2021). We sought to explore roles for neurexins in dense core vesicle and neuropeptide release using the common coelomocyte uptake assay in *C. elegans* (Sieburth, Madison, and Kaplan 2007; Sasidharan et al. 2012; Gracheva et al. 2007; Tikiyani et al. 2018; Babu et al. 2011; Pandey, Bhardwaj, and Babu 2017). The *nlp-21* neuropeptide gene is broadly expressed in cholinergic and GABA neurons (CeNGEN (Hammarlund et al. 2018), and **Supplemental Table 1**)). We analyzed the levels of a fluorescently tagged NLP-21 neuropeptide transgene expressed in cholinergic motor neurons (*unc-129*p*::nlp-21::Venus*). We measured levels of expression in dorsal nerve cords (unsecreted fraction) and coelomocytes (secreted fraction) in controls and loss of function *nrx-1* alleles (**Figure 1A** (adapted from (Gracheva et al. 2007)). In *C. elegans*, the single *nrx-1* gene encodes 2 major isoforms (a long α and a short γ) that have neuron and synapse specific roles (Kurshan et al. 2018; Bastien, Cowen, and Hart 2023).

**Figure 1.**
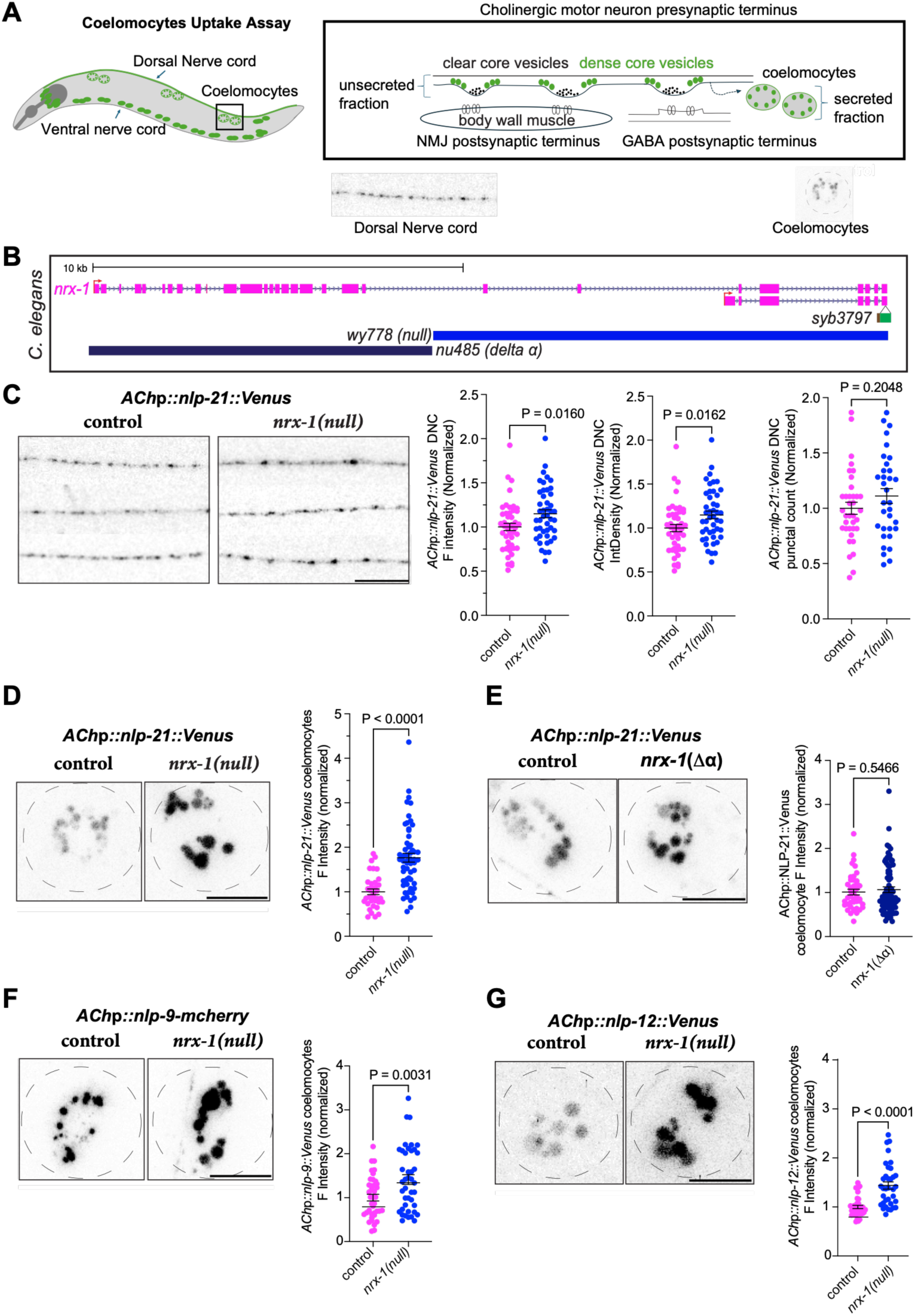
*nrx-1* regulates the secretion of multiple NLP family neuropeptides from cholinergic motor neurons. **A)** Schematic of Coelomocyte uptake assay, **B)** Schematic of the *nrx-1* gene and alleles used in this study. **C**) representative confocal micrographs of dorsal nerve cords of controls and *nrx-1(null*) mutants expressing NLP-21::Venus in cholinergic motor neurons and graphs showing quantification of normalized florescence intensity, integrated density, and punctal count in both genotypes. Representative confocal micrographs and graphs quantifying normalized florescence intensity of coelomocytes for **D)** cholinergic NLP-21::Venus (*unc-129*p), **E)** *α*-isoform specific deletion of *nrx-1(nu485*), **F)** cholinergic NLP-9::mcherry (*acr-2*p), and **G)** NLP-12::Venus (*unc129*p). Scale bar=10µm.

**Supplemental Figure 1.**
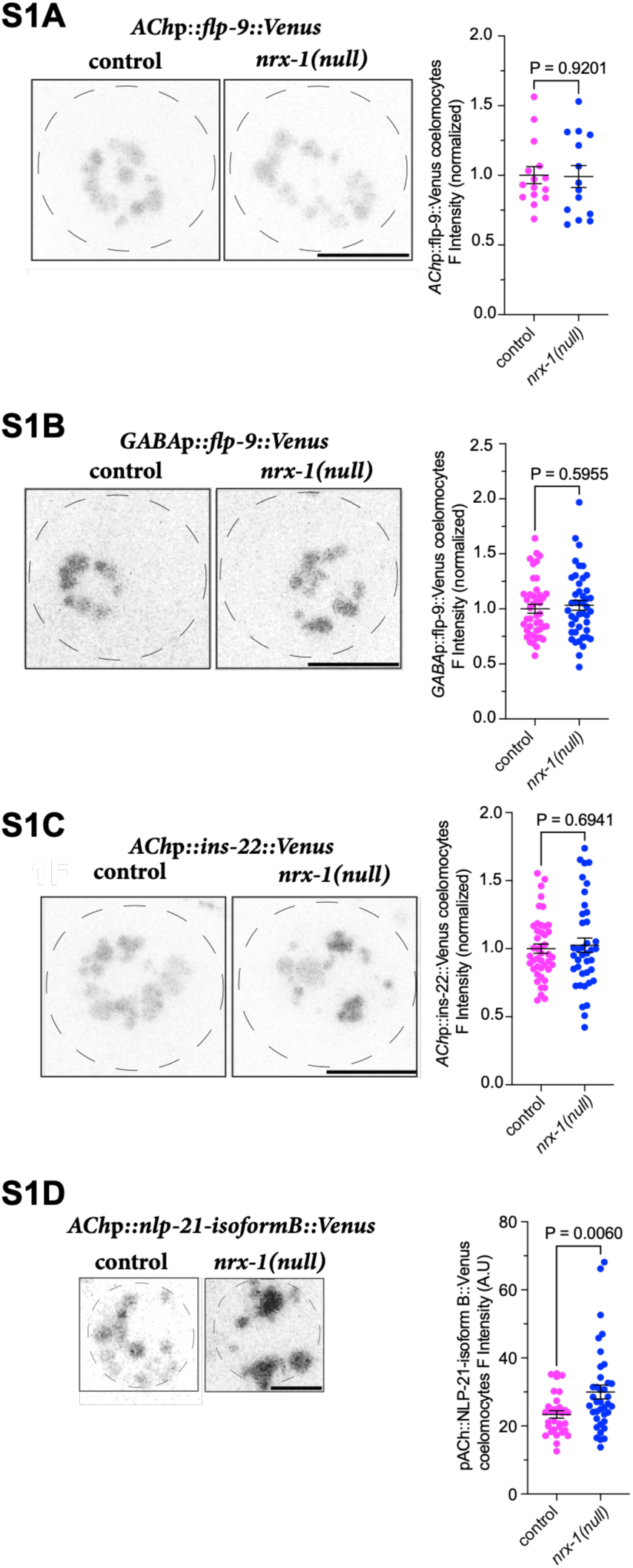
n*r*x*-1* does not impact release of INS-22 or FLP-9 neuropeptides. A and. **B)** representative coelomocyte images and normalized graphs of control and *nrx-1(wy778*) mutants expressing FLP-9::Venus either in cholinergic or GABA neurons, **C)** representative images and normalized graph cholinergic INS-22::Venus (*unc-17*p) in controls and *nrx-1(wy778*) mutants S1D) representative images and normalized graph for control and *nrx-1(wy778*) mutants expressing ß-isoform of NLP-21::Venus under cholinergic promoter *unc-129* (extrachromosomal array).

We tested a null allele *wy778* that deletes both α- and γ-isoforms and an α-isoform specific allele *nu485* (**Figure 1B**). We observed that NLP-21::Venus was localized in a punctate pattern along the dorsal nerve cord as previously described in controls and animals lacking *nrx-1(wy778)*(**Figure 1C**). We found a slight increase in levels of NLP-21::Venus in *nrx-1* null animals in the dorsal nerve cords compared to controls (**Figure 1C**). After release or secretion from the cholinergic motor neurons NLP-21::Venus gets endocytosed by the coelomocytes, which can be measured to quantify levels of released neuropeptide. We found that NLP-21::Venus in the coelomocytes was increased in *nrx-1(wy778)* null animals compared to control animals (**Figure 1D**). Since the coelomocyte secretion was a more robust phenotype, we focus our further experiments on this phenotype. To explore the *nlp-21* neuropeptide phenotype, we generated additional *nlp-21* expression constructs expressing α- and β-isoforms of *nlp-21* under a cholinergic neuron promoter (*unc-129*p), tagged with Venus reporter. We observed expression of β-isoform NLP-21::Venus similar to the integrated transgene (*nuIs183*: *unc129*p*::nlp-21::Venus*) and observed a similar increase in coelomocyte fluorescence levels in *nrx-1* null animals compared to controls (**Supplemental Figure 1D**). However, when we tried to express the α-isoform of NLP-21::Venus, we did not see any expression even when injected at higher concentration (80-100ng/ul)(data not shown). Together, these results indicate that loss of neurexin slightly increases localization of NLP-21::Venus in the dorsal nerve cord and increases secretion of NLP-21::Venus from cholinergic motor neurons.

We next asked if *nlp-21* secretion was impacted in α-isoform specific *nrx-1* alleles and found no change in NLP-21::Venus secretion compared to controls with a large deletion of the α-isoform (*nu485*)(**Figure 1E**). This indicates that loss of both isoforms is required to increase NLP-21 secretion, and that the long α isoform is dispensable for regulating this phenotype. Next, we asked if secretion of other neuropeptides is also altered in *nrx-1* null mutants. In *C. elegans*, 113 neuropeptide genes encoding over 250 distinct neuropeptides have been identified which are classified into 3 groups: 1) insulin-like peptides (INS), 2) FMRFamide-related peptides (FLPs), and non-insulin, non-FMRFamide-related neuropeptides (NLPs)(Li and Kim 2010). We tested INS-22::Venus expressed in cholinergic neurons and found no change in the coelomocyte uptake of this peptide in *nrx-1* null mutants compared to control animals (**Supplemental Figure 1C**). We also tested expression of FLP-9::Venus, expressed either in cholinergic or GABA motor neurons, and again found no change in coelomocyte florescence of this peptide in the *nrx-1* null mutants compared to controls (**Supplemental Figure 1B and 1C**, respectively). We tested 2 more neuropeptides from the nlp group (NLPs), NLP-9::Venus and NLP-12::Venus expressed in the cholinergic neurons and found a similar increase in the secretion levels of these neuropeptides in the *nrx-1* null animals compared to controls (**Figure 1F&G**). This indicates that NRX-1 regulates secretion of multiple nlp neuropeptides from cholinergic motor neurons.

### *nrx-1* functions cell-autonomously in cholinergic motor neurons to regulate NLP-21 secretion

At the *C. elegans* cholinergic neuromuscular junctions (NMJs), NRX-1 was found to function in the postsynaptic muscles through an interaction with presynaptic NLG-1, where it inhibits release of acetylcholine neurotransmitters via a retrograde signaling (Hu et al. 2012). However at dyadic synapses from cholinergic motor neurons onto postsynaptic GABAergic motor neurons and muscle cells, NRX-1 is localized pre-synaptically, where it regulates ACR-12 (acetylcholine receptor) clustering in postsynaptic GABAergic neurons, but not in muscles (Philbrook et al. 2018). Further, at the GABAergic NMJs, NRX-1 is presynaptic and regulates postsynaptic GABA receptor clustering via its interaction with postsynaptic NLG-1 and presynaptic partner, MADD-4 (Maro et al. 2015). The pleiotropic function and localization of NRX-1 led us to test if NRX-1 is functioning at the presynaptic or postsynaptic sites to regulate NLP-21::Venus secretion.

To localize NRX-1 function, we knocked down the endogenous NRX-1 in a spatiotemporal manner using the Auxin Inducible Degron (AID) system (**Figure 2A&B**). Briefly, a super folder GFP and the AID tag (sfGFP::AID::linker) were integrated into the 3’ end of the endogenous *nrx-1* locus and the E3 ubiquitin ligase for AID, TIR1, was expressed either in neurons (*rgef-1* promoter (Bastien, Cowen, and Hart 2023)) or in muscles (*myo-3* promoter (Ashley et al. 2021)). Animals expressing both NRX-1::sfGFP::AID and neuronal or muscle TIR1 were grown with and without auxin for at least 2 generations to maximize the knockdown of NRX-1 throughout the developmental and adult periods. We found that neuronal expression of TIR1 with auxin increased the NLP-21::Venus fluorescence intensity in coelomocytes compared to controls with no auxin (**Figure 2C**), similar to what we observed with the *nrx-1* null mutant allele (**Figure 1D**). We did not detect any changes in the levels of NLP-21::Venus secretion with expression of TIR1 in the muscles (**Figure 2C**), indicating that NRX-1 is acting in neurons and not in muscles for the release of NLP-21::Venus. We also checked if auxin treatment changes NLP-21::Venus secretion, but found that 1 mM auxin had no impact in the control strain (*unc129*p*::nlp-21::Venus* (**Supplemental Figure 2A**)).

**Figure 2.**
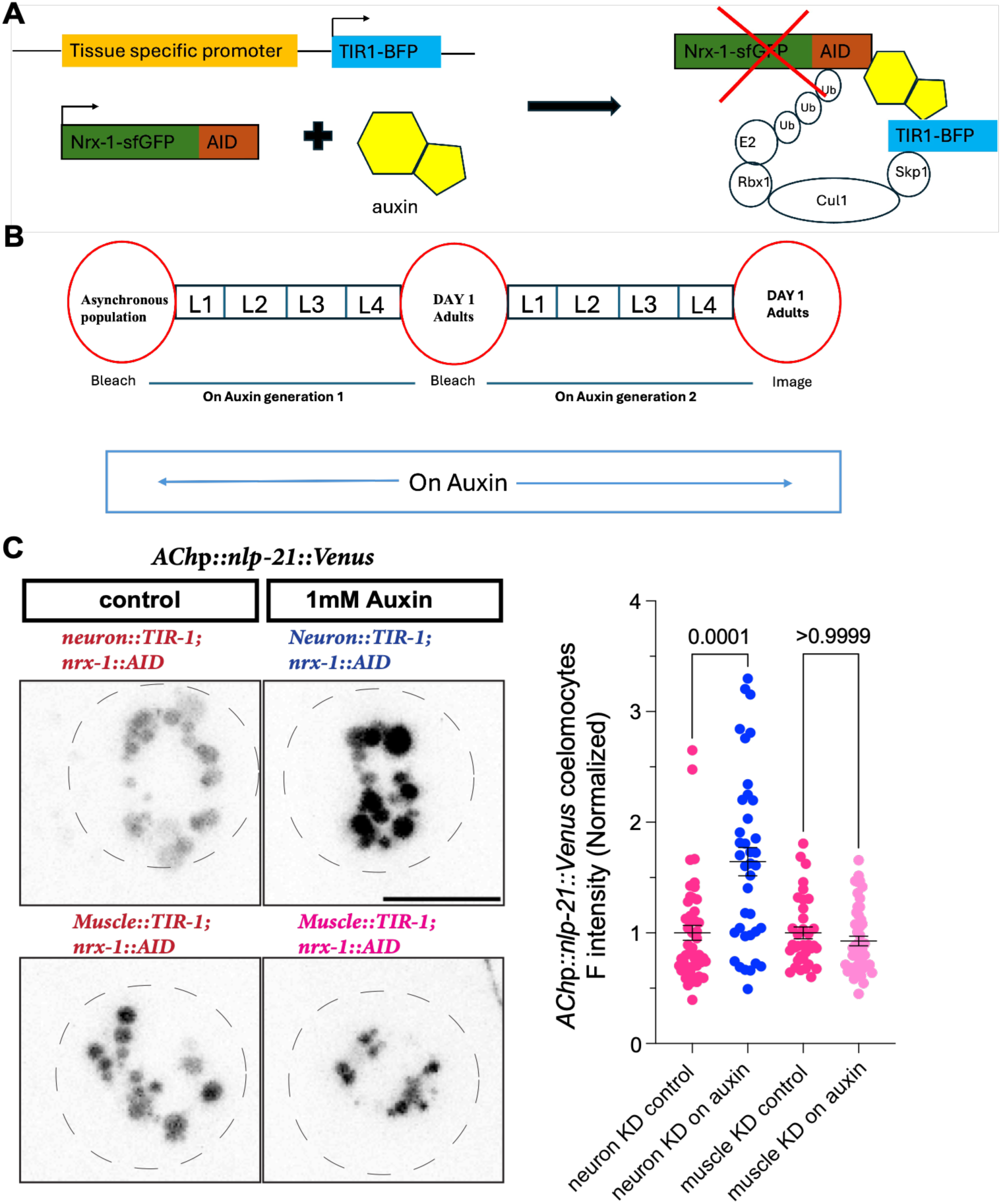
*nrx-1* functions in neurons to regulate the secretion of NLP-21 neuropeptide. **A)** Schematic of Auxin-Inducible Degron (AID) system used to degrade endogenous NRX-1 in a tissue-specific manner-TIR1 tagged with BFP reporter protein is expressed either pan-neuronally using *rgef-1* promoter or in the muscles using *myo-3* promoter. NRX-1::sfGFP with AID (degron) was integrated in the genome using a CRISPR knockin. **B)** Experimental strategy used to degrade NRX-1 in a temporal manner. Bleach synchronized worms were put on 1mM auxin for at least 2 generations to completely degrade NRX-1 throughout the development including in embryos and eggs. **C)** Representative confocal micrographs of coelomocytes in animals expressing NRX-1::AID and TIR-1 (either under neuronal promoter or muscle promoter) without (control) and with Auxin (1mM) expressing cholinergic NLP-21::Venus and the normalized florescence intensity graph.

**Supplemental Figure 2.**
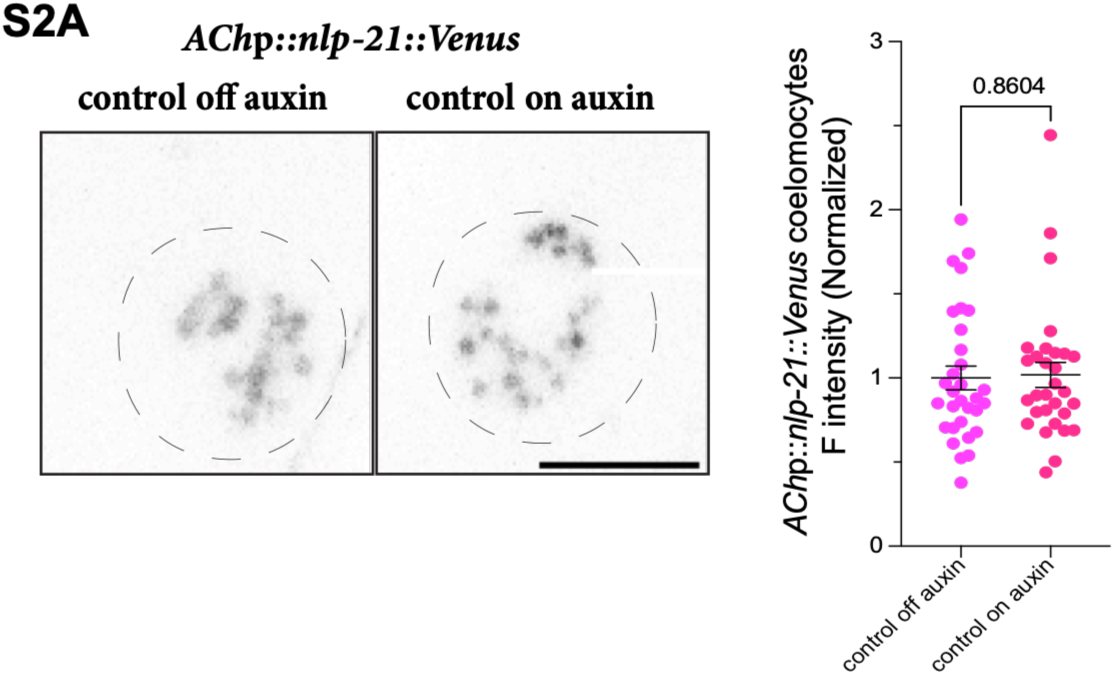
Auxin exposure alone does not impact NLP-21 release. Representative confocal micrographs of coelomocytes and graph of NLP-21::Venus without and with 1mM auxin treatment.

We next extended this result to test which NRX-1 isoform is required in neurons and cell-autonomously in the cholinergic neurons to regulate the release of NLP-21. We expressed either the long α-isoform or short γ-isoform of NRX-1 (protein domain structures are shown in **Figure 3A**) in all neurons using a neuronal promoter (*ric-19*) in *nrx-1* null mutants (*wy778*) and found that both isoforms decreased NLP-21::Venus secretion in the coelomocyte uptake assay compared to *nrx-1(wy778)* mutants alone, and similar to control animals (**Figure 3B**). This indicates that NRX-1 is required in the neurons to regulate secretion, and that both isoforms when overexpressed in neurons are sufficient for this function. Next, we expressed the long α-isoform of NRX-1 in muscles using the *myo-3* promoter. Muscle expression failed to rescue the NLP-21::Venus secretion phenotype of *nrx-1* null mutants, indicating that NRX-1α cannot function in muscles to regulate NLP-21 secretion. Next, we expressed NRX-1 in cholinergic (*unc129*p) or GABAergic (*unc-47*p) motor neurons using the same strategy and found that expression of NRX-1 isoforms in cholinergic neurons, but not GABAergic neurons, decreased NLP-21::Venus secretion compared to *nrx-1(wy778)* mutants alone (**Figure 3C&D**). At the cholinergic NMJs, NRX-1 binds to NLG-1 to regulate acetylcholine release (Hu et al. 2012), we wondered if NLG-1 is also required to maintain NLP-21secretion, we checked cholinergic NLP-21 secretion in *nlg-1(ok259)* mutants and found that *nlg-1* mutants show NLP-21 levels similar to controls (**Supplemental Figure 3A**), indicating NRX-1 either does not require a ligand to regulate NLP-21 secretion or it is binding to a noncanonical partner to maintain this function. Taken together, these experiments indicate that NRX-1 functions cell autonomously in the cholinergic motor neurons to regulate release of NLP-21, and that both isoforms can rescue the loss of function phenotype when overexpressed in the same neurons.

**Figure 3.**
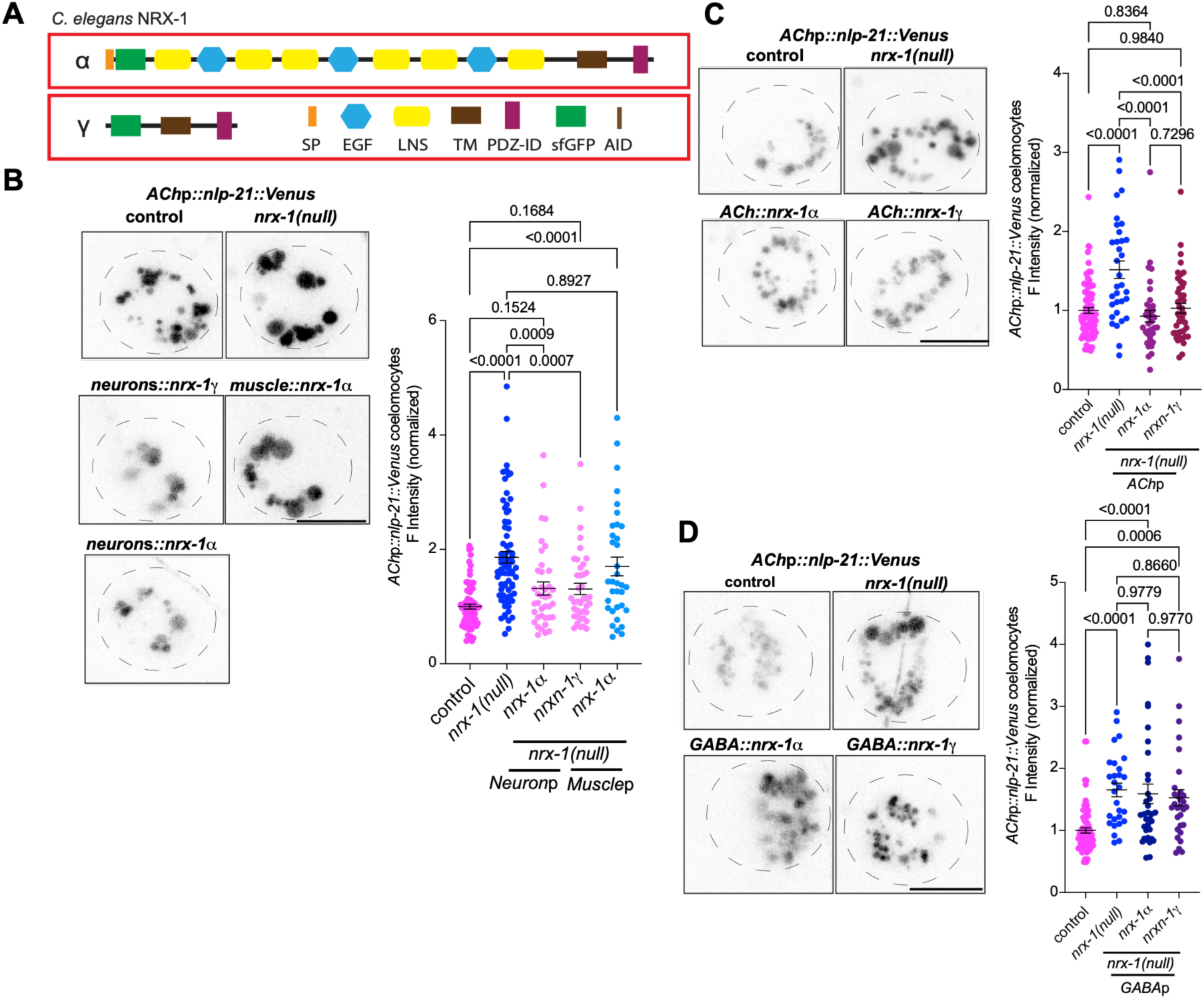
*nrx-1* functions cell-autonomously in cholinergic motor neurons to regulate NLP-21 secretion. **A)** A cartoon showing the protein domains of the long isoform of nrx-1(α) and the short isoform of nrx-1(γ). **B)** Representative confocal micrographs of coelomocytes in controls and *nrx-1(wy778)* mutants expressing cholinergic NLP-21::Venus and the normalized florescence intensity graph. NRX-1 isoforms were expressed via extrachromosomal arrays with a neuronal promoter (*ric-19*p*::sfGFP::nrx-1α* or *ric-19*p*::sfGFP::nrx-1*γ), or muscle promoter (*myo-3*p*::sfGFP::nrx-1α*). **C)** and **D)** Representative confocal micrographs of coelomocytes of controls and *nrx-1(wy778*) mutants expressing NLP-21::Venus and the normalized florescence intensity graph. The long isoform of nrx-1(α) and the short isoform of nrx-1(γ) were expressed via extra chromosomal array under the cholinergic promoter (*unc-129*p) or the GABAergic promoter (*unc-47*p), respectively.

**Supplemental Figure 3.**
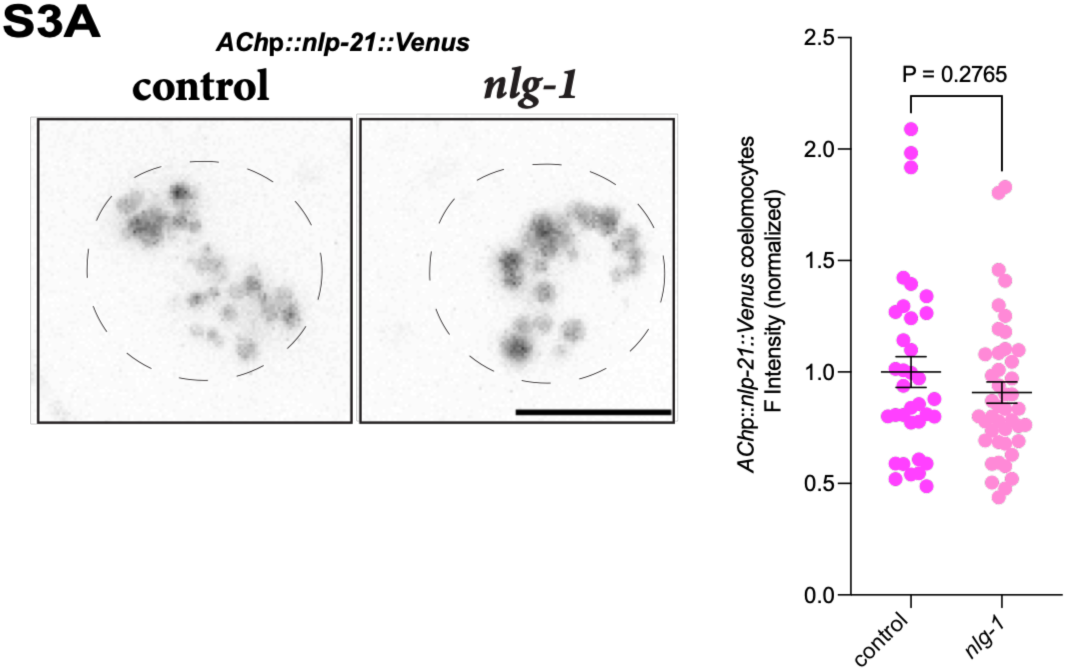
n*r*x*-1’s* canonical binding partner *nlg-1* is not required to maintain NLP-21 secretion. Coelomocytes florescence intensity and quantification graph for the control and *nlg-1(ok259)* mutants expressing *unc129*p*::nlp-21::Venus* transgene.

### *nrx-1* regulates dense core vesicle and active zone morphology in cholinergic motor neurons

We next asked how NRX-1 is regulating the release of NLP-21 neuropeptide, whether it is acting as a scaffolding protein to keep the dense core vesicles at the synapses or is involved in pathways responsible for the release of these neuropeptides. To address the first possibility, we first analyzed NRX-1 localization and potential to interact with dense-core vesicles. We tagged the dense-core vesicle marker protein, IDA-1 (PTPRN)(Zahn et al. 2004) and expressed it in cholinergic motor neurons (*unc-129*p*::ida-1::mRuby*) in a strain with endogenously tagged *nrx-1* (*nrx-1::sfGFP::AID*). We observed a punctate pattern of expression along the dorsal nerve cords for both the proteins and that some of the IDA-1 puncta in cholinergic motor neurons was colocalized with NRX-1 (**Figure 4A**). Next, we asked about the percentage of dense core vesicles carrying NLP-21 neuropeptide, since NLP-21 should be packaged in the dense core vesicles. We crossed NLP-21::Venus with IDA-1::mRuby (both cholinergic promoters) and observed clear colocalization of both proteins, indicating that NLP-21 is packaged into and included in most of the cholinergic dense core vesicles (**Supplemental Figure 4D**). we then checked the localization of NLP-21 and IDA-1 in *nrx-1 (wy778)* animals and observed that although NLP-21 puncta still colocalize with IDA-1, but the levels of IDA-1 were reduced in *nrx-1(wy778)* compared to controls (**Supplemental Figure 4D**).

**Figure 4.**
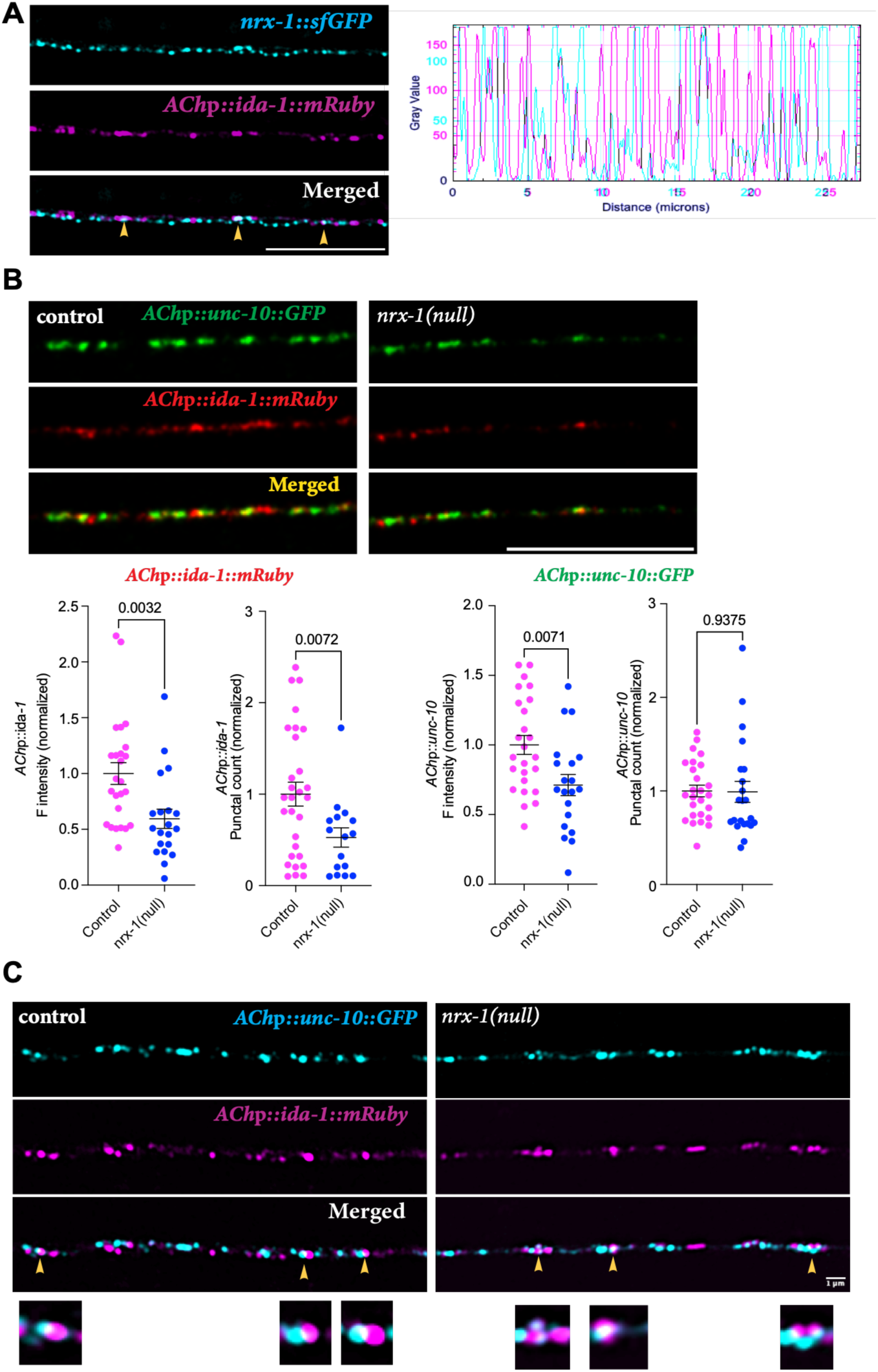
Loss of *nrx-1* disrupts dense core vesicle expression and localization in cholinergic motor neurons. **A)** representative enhanced resolution confocal micrographs of dorsal nerve cords of NRX-1::sfGFP::AID and dense core vesicle marker, IDA-1::mRuby under cholinergic promoter (*unc-129*p). Yellow arrows show partial colocalization of NRX-1 puncta with IDA-1 puncta, and the graph shows fluorescence intensity peak across the dorsal nerve cord axis **B)** representative confocal micrographs of dorsal nerve cords expressing active zone protein, UNC-10 and dense-core vesicles protein, IDA-1 in cholinergic motor neurons (*unc-129*p) in controls and *nrx-1(wy778*) mutants. Quantitation graphs showing normalized fluorescence intensity and number of puncta for IDA-1 and UNC-10, respectively (Scale bar=10µm, Statistics: Student’s t-test and P<0.05). **C)** Representative Airyscan confocal micrographs of dorsal nerve cords expressing active zone protein UNC-10 and dense-core vesicles protein, IDA-1 in cholinergic motor neurons (*unc-129*p) in controls and *nrx-1(wy778*) mutants. Yellow arrowheads show juxtaposition of the puncta.

**Supplemental Figure 4.**
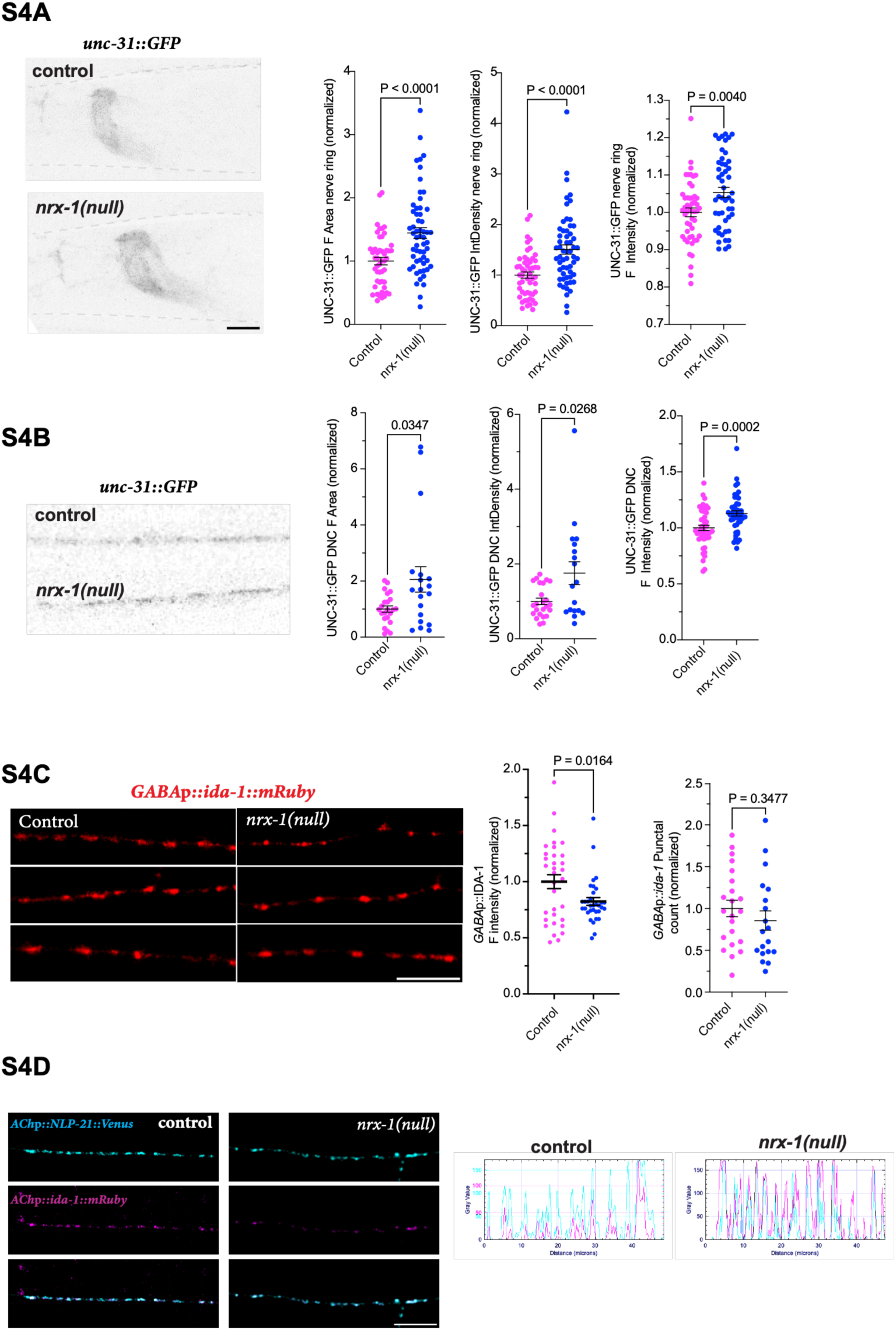
Loss of *nrx-1* alters UNC-31 levels and reduces dense core vesicle markers in GABAergic motor neurons. **A)** and **B)** Representative images of the head regions of control and *nrx-1(null)* animals showing the expression of UNC-31::GFP in the nerve ring and dorsal nerve cords and the quantification graphs showing UNC-31 normalized fluorescence area, integrated density, and fluorescence intensity, (Scale bar=10µm). **C)** Representative images of dorsal nerve cords of control and *nrx-1(wy778*) worms expressing IDA-1::mRuby in GABA neurons (*unc-47*p) and the quantitation of IDA-1 fluorescence intensity and punctal count normalized to control. scale bar=10µm. **D)** Representative dorsal nerve cord images of control and *nrx-1 (null)* animals expressing NLP-21 and IDA-1 in cholinergic neurons, and the graphs show relative fluorescence peaks for both the puncta across the cord in both the genotypes. scale bar=10µm.

We then analyzed the morphology of cholinergic motor neuron synapses by analyzing the active-zone marker protein, UNC-10 (RIMS), together with dense-core vesicles marker, IDA-1 and performed co-localization experiments. We observed limited colocalization between these proteins (**Figure 4B**), indicating the dense-core vesicles release sites are largely distinct from the active zone release sites, which may be considered extra synaptic. When we looked at these proteins together in *nrx-1(wy778)* null animals, we observed that UNC-10 protein levels were decreased similar to a previous report for active zone marker, *cla-1::GFP* (Kurshan et al. 2018). Remarkably, we observed that for dense-core vesicle marker, IDA-1, both puncta fluorescence intensity and puncta count were also decreased in *nrx-1* animals compared to controls (**Figure 4B**).

To further analyze the localization and morphology of these two proteins and synaptic morphologies, we used enhanced-resolution confocal microscopy. We visualized UNC-10 and IDA-1 in controls and *nrx-1*(null) animals and observed that both proteins were juxtaposed to each other with a clear round shaped morphology in controls. However, in *nrx-1* null mutants, the localization and morphology of these proteins were disrupted, with the proteins appearing somewhat merged or overlapping compared with controls, where a clear boundary or spatial segregation between the proteins is maintained (**Figure 4C**). We also analyzed IDA-1 expression and localization in the GABA motor neurons, and observed a significant decrease in the fluorescence intensity, but no change in the punctal count of dense-core vesicles in *nrx-1* null mutants (**Supplemental Figure 4C**), indicating that NRX-1 can regulate the localization of DCVs in both cholinergic and GABAergic motor neurons. We next tested expression of the DCV priming factor and neuropeptide release regulator, UNC-31/CADPS. Endogenous UNC-31::GFP fluorescent area, integrated density, and fluorescence intensity were increased in the nerve ring and dorsal nerve cords of *nrx-1* null animals (**Supplemental Figure 4A and 4B**). Taken together, these results indicate that NRX-1 regulates the clustering, distribution and expression of the DCV protein IDA-1 (PTPRN), maintains separation and juxtaposition of neurotransmitter and neuropeptide release sites, and can regulate the expression of DCV secretion regulator UNC-31 (CADPS). The increase in a priming and release regulator with loss of *nrx-1* could mediate the increase in NP release we observe, although not the NP-specificity of this phenotype.

### NLP-21 release is not impacted by excitation/inhibition disruption, but is dependent on Ca²⁺ channel function

Maintaining proper balance of excitation and inhibition (E/I) is critical for neuronal function and to maintain brain homeostasis. *nrx-1* is crucial for cholinergic and GABAergic motor neuron synapse function and excitation/inhibition (E/I) balance at neuromuscular synapses (Hu et al. 2012). We asked whether disruption of cholinergic excitation (*unc-17*) or GABAergic inhibition (*unc-25*) impacts NLP-21 release, but found no change in cholinergic NP release in either mutant (**Figure 5A**), suggesting that the increased NLP-21 release with loss of *nrx-1* is not solely due to disrupted synaptic activity or E/I imbalance. Neurotransmitter release from clear core vesicles and neuropeptide release from dense-core vesicles are strongly Ca²⁺ dependent, with increased intracellular Ca²⁺ influx enhancing DCV exocytosis and neuropeptide secretion (Kreutzberger et al. 2017; Engisch and Nowycky 1996; Xia, Lessmann, and Martin 2009). To test if changes in calcium levels impact NLP-21 secretion, we tested NLP-21 secretion in *unc-2 (lof)* mutants. UNC-2 is the *C. elegans* homolog of the CaV2 voltage-gated calcium channel α1 subunit (closely related to mammalian CaV2.1/CACNA1A and CaV2.2/CACNA1B). It is a major presynaptic calcium channel that triggers neurotransmitter release and has been shown to be regulated by NRX-1/NLG-1 signaling at the *C. elegans* cholinergic NMJs (Hu et al. 2012). *unc-2* mutants show a reduction in NLP-21 secretion (**Figure 5B**) while *unc-2;nrx-1(wy778)* double mutants suppressed the *nrx-1(wy778)* mutant phenotype of increased NLP-21 secretion, with NLP-21 secretion comparable to *unc-2* single mutants (**Figure 5B**). This suggests that the increase in NP release in animals lacking *nrx-1* is dependent on *unc-2* and that decreased intracellular levels in the absence of *unc-2* function leads to reduced NLP-21 secretion. These results indicate that the *nrx-1* NLP-21 release phenotype is not simply a consequence of altered excitation/inhibition that could result from other functions of *nrx-1*, but may reflect dysregulation of pre-synaptic intracellular Ca²⁺-dependent mechanisms that control neuropeptide secretion, and may also explain the lack of phenotype in *nlg-1* mutants.

**Figure 5.**
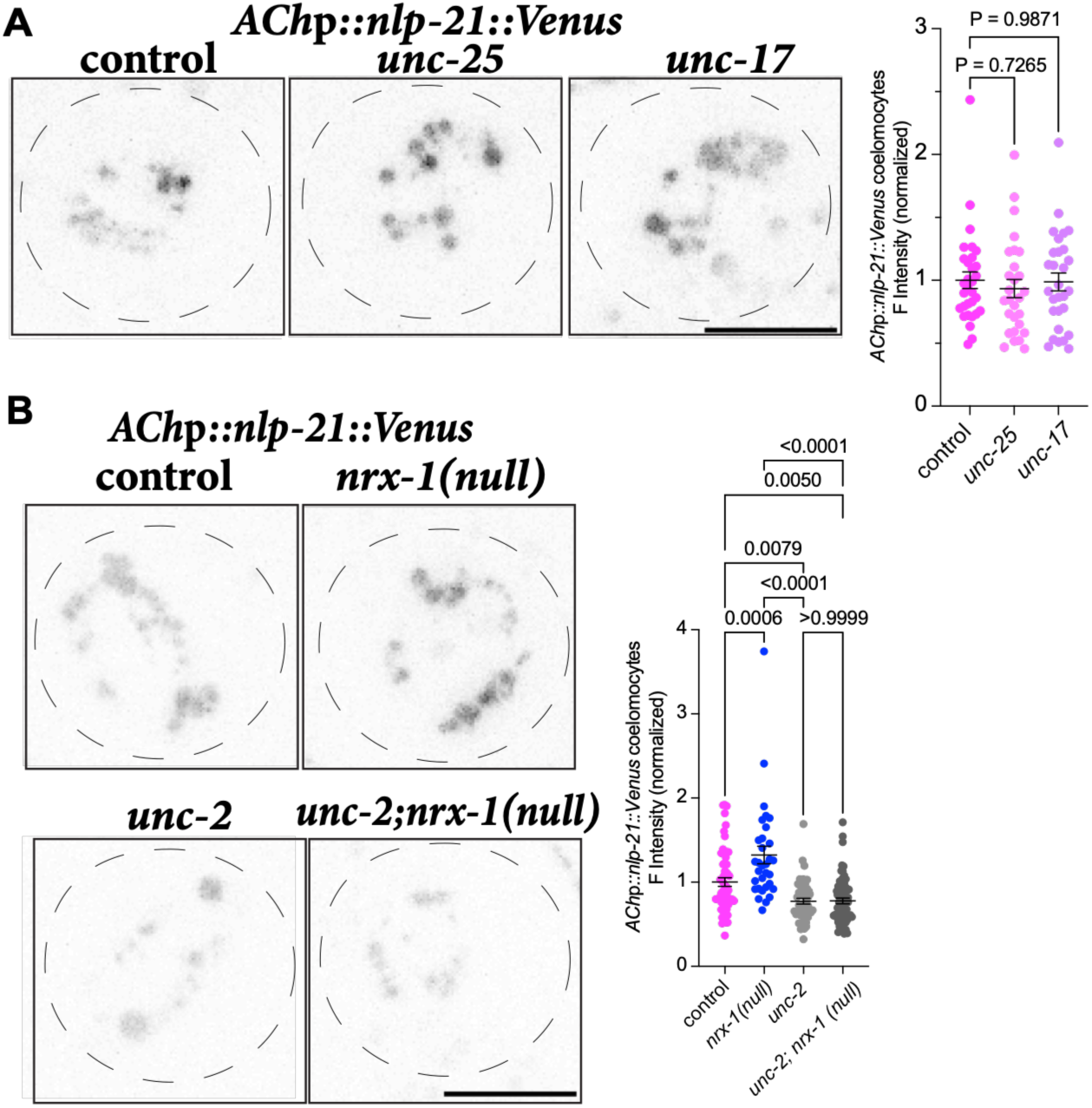
NLP-21 release is independent of alterations in excitation/inhibition but is regulated by intracellular Ca²⁺ levels. Representative images of coelomocytes of **A)** control, *unc-25(e156)*, and *unc-17 (e245)* mutants and **B)** control, *nrx-1 (null)*, *unc-2 (gk366), and unc-2 (gk366);wy778* expressing NLP-21::Venus in cholinergic neurons and their fluorescence intensity measurement graph. scale bar=10µm.

### A rare autism variant in *nrx-1* results in a gain of function NLP-21 secretion phenotype

To test the potential relevance of NRX-1 and NP signaling to human genetics, we checked if cholinergic NLP-21 secretion is impacted by insertion of a rare variant found in an autism proband. We have found that this variant impacts multiple behaviors in *C. elegans* and has gain of function phenotypes compared to loss of *nrx-1* null allele (Haskell et al. 2025). The rare variant allele, *nrx-1*(s*yb869*), changes a leucine to glutamine at the 16th amino-acid (L16Q) in the first exon (**Figure 6A**)(Haskell et al. 2025). The *nrx-1(syb869)* mutants have reduced NLP-21 secretion compared to control animals (**Figure 6B**), an opposite phenotype to that observed in *nrx-1* null animals (**Figure 1D**), and have no change in the unsecreted fraction in the dorsal nerve cord (**Supplemental Figure 6A**). This finding has interesting implications for our recent finding that the *nrx-1(syb869)* allele induces novel gain-of-function phenotypes in solitary feeding and activity levels in response to food and loss of food (Haskell et al. 2025). These results indicate that neurexin’s role may not be a simple on/off switch for excitation/inhibition but that variation in the nrx-1 gene and protein alter neurons and circuits in different directions, which could explain why *NRXN1* variants in humans produce heterogeneous clinical presentations across the autism spectrum, rather than one uniform “hyperexcitable” or “hypoexcitable” phenotype.

**Figure 6.**
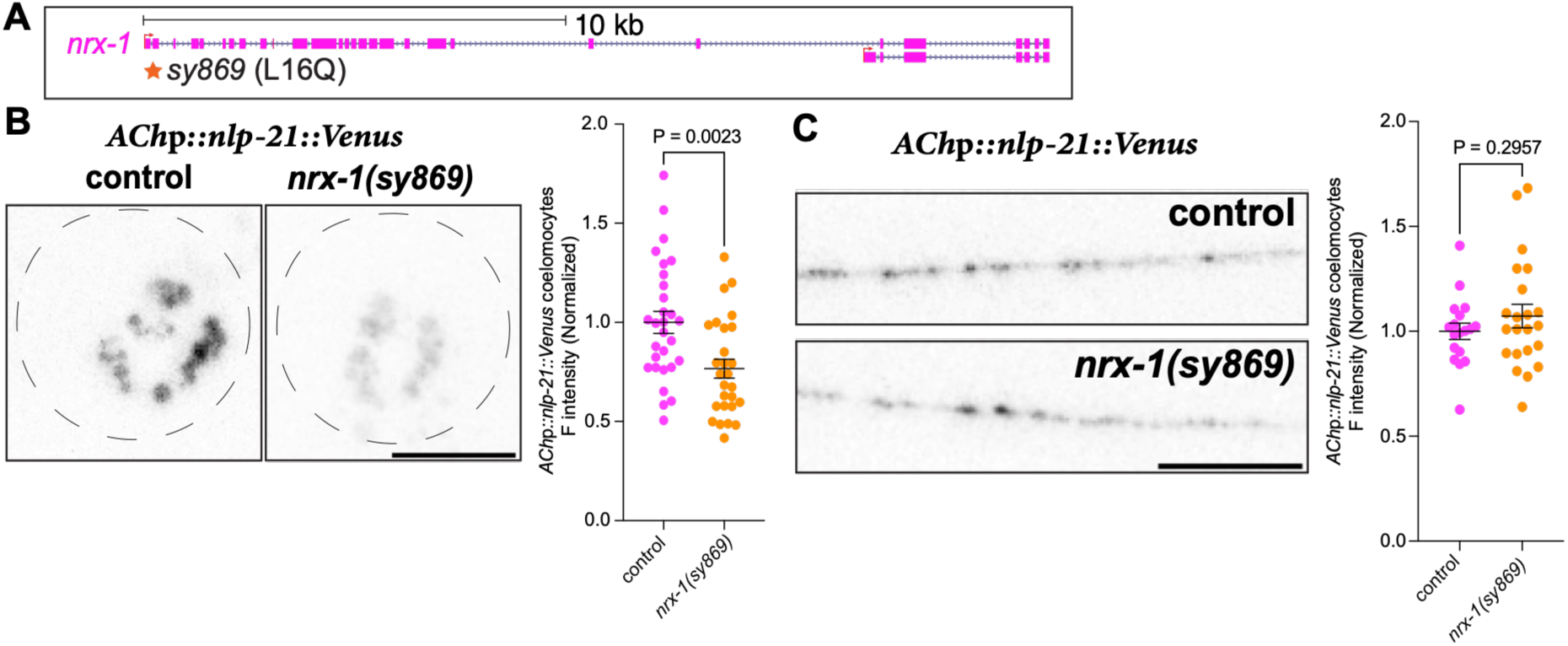
Conserved autism allele of NRX-1(L16Q) reduces NLP-21 secretion. **A)** Schematic of *nrx-1* gene showing the location of *sy869* allele **B) and C)** representative images and the quantification graphs for the coelomocytes and the dorsal nerve cords of the control and *nrx-1(syb869)* animals expressing *unc-129*p*::nlp-21::Venus*. Scale bar-10µm.

## DISCUSSION

Here, we report that *nrx-1* regulates the release of multiple neuropeptides from cholinergic motor neurons in *C. elegans*. Using tissue specific transgenic expression and neuron-specific degradation of endogenous NRX-1 protein, we show that *nrx-1* functions in a cell-autonomous manner. We find that isoforms of *nrx-1* are restrict NP secretion from the cholinergic neurons. We confirm that loss of the *nrx-1* impacts cholinergic active-zone number and morphology, but also find it impacts the clustering, distribution, and expression of the DCV protein, IDA-1 (PTPRN), and DCV secretion regulator, UNC-31 (CADPS). Our results suggest that *nrx-1* acts as to maintain separation and juxtaposition of neurotransmitter and NP release sites and impacts DCV localization. *nrx-1* is crucial for cholinergic and GABAergic motor neuron synapse function and excitation/inhibition (E/I) balance at neuromuscular synapses, but we find that loss of cholinergic excitation (*unc-17*) or GABAergic inhibition (*unc-25*) does not alter release of cholinergic NPs, suggesting that increased release upon loss of *nrx-1* is not solely due to disrupted synaptic activity or E/I imbalance.

Although SAMs are well established regulators of synaptic vesicle release (reviewed in (Gottmann 2008; Luo et al. 2020; Missler 2003)), emerging evidence suggests that they may also influence neuropeptide-containing DCVs. Ferdos et al. showed that deletion of all three β-neurexin isoforms reduced the number of presynaptic DCVs and chromogranin A-positive vesicles in cultured mouse hippocampal neurons and cerebellar tissue. However, the study did not directly measure DCV exocytosis or neuropeptide release, leaving open the question of whether neurexins regulate neuropeptide secretion itself (Ferdos et al. 2021). At the synapse, the neurexin superfamily can interreact with more than 50 different ligands (Boucard, Ko, and Südhof 2012), and one of the transsynaptic binding partners, latrophilins controls the release of insulin hormone containing large dense-core vesicles from the pancreatic primary β-cells and INS-1/MIN6 lines in the presence of α-latrotoxin (Lang et al. 1998). More recently, it has been shown that latrophilins are highly expressed across islet cell types, and distinct LPHN3/ADGRL3 splice variants reduce insulin secretion from β-cells by lowering cAMP via Gi-mediated pathway (Röthe et al. 2019). Further, neuroligins, neurexins, and SynCAMs are expressed in β-cells, and neuroligin 2 overexpression and knockdown altered insulin secretion in INS-1 cells (Suckow et al. 2008). Clustered, but not soluble neuroligin 2, increased insulin secretory capacity, and the effect occurred distal to glucose sensing, while neurexin binding was not required (JBC 2012). In neuroligin 2 knockout mice, insulin granule docking was reduced while secretion was paradoxically increased, with lower neurexin transcript levels (PLoS One 2013). There is an inconsistency in the directionality across acute and chronic manipulations, which may be due to compensation.

A *C. elegans* study has recently implicated the conserved synaptic adhesion protein CASY-1/calsyntenin in neuropeptide signaling. The study reports reduced FLP-21 accumulation/release in *casy-1* mutants; however, whether this phenotype reflects impaired DCV transport or exocytosis remains unresolved (Shahi, Thapliyal, and Babu 2025). However, no adhesion molecule has been shown to organize DCV fusion sites. Release sites and organizing principles for DCV exocytosis remain elusive, and how the shared SNARE machinery is transiently assembled at DCV fusion sites is unresolved (Moro et al., *Sci Adv* 2021). Here, we show that a SAM, *nrx-1,* regulates the release of specific subset of neuropeptides from cholinergic neuromuscular junctions (NMJs) in *C. elegans* by organizing the distribution and localization of DCVs.

Of the neuropeptides examined, three belonging to the NLP class showed increased release, while two others from different classes remained unchanged. It will be interesting to determine why and how *nrx-1* discriminates between these neuropeptide classes, and how their release mechanisms differ. Specifically, whether all classes depend on the same calcium-dependent SNARE machinery, with *nrx-1* acting selectively on NLP-class release, or whether distinct release machineries govern the other classes. Because UNC-31/CADPS is required for dense-core vesicle release generally, a global increase in priming factor would be expected to elevate secretion of all DCV cargo. We instead observe increased release of three NLP-class peptides with no change in INS-22 or FLP-9, and no change in UNC-31 levels in the nerve cord. Peptide-class selectivity is therefore difficult to explain by priming factor abundance alone, and instead favors a mechanism involving spatial reorganization of release sites or differential regulation of DCV subpopulations. Additionally, given the many behavioral phenotypes observed with loss of *nrx-1*, whether it functions in different neuron types to regulate different NPs may have large implications for these behavioral phenotypes. To further understand the mechanisms by which *nrx-1* is regulating NP release, we are currently testing if genes known to regulate release of NPs including *ric-7, tom-1* (TOMOSYN), and *pkc-1* (PRKCE) are involved in the NP release phenotype we observe. We are also analyzing behavioral phenotypes of the neuropeptides and *nrx-1* genes to identify functional impact of this phenotype, and whether loss of neurexins regulates NP release and DCV morphology in mammalian neurons. We find that neurexins regulate NP signaling, a significant novel function for neurexin genes in regulating circuits and behaviors, and of potential importance for associated human conditions.

### Implications for humans *NRXN1* associated disorders and conditions

Disruption of human *NRXN1, NRXN2*, and *NRXN3* genes, including deletions and copy number variations, are associated with autism spectrum, schizophrenia, intellectual disability, and ADHD (AlHousni et al. 2026; Selen et al. 2026; Friedman et al. 2026; Liu et al. 2026; Tromp, Mowry, and Giacomotto 2021; Zhong et al. 2022). *NRXN* gene disruptions can cause variable penetrance, genetic-background effects, pleiotropy, and circuit-specific effects to shape these diverse behavioral phenotypes, rather than producing a simple one-to-one genotype–disease relationship (Fuccillo and Pak 2021; Khoja, Haile, and Chen 2023; Cameli et al. 2021; Al Shehhi et al. 2019; Gomez, Traunmüller, and Scheiffele 2021; Cao and Tabuchi 2017; Fernando et al. 2025). How disruption of neurexin genes, even by deletions or variants with similar predicted impact on gene function, can result in such heterogeneous behavioral changes is an open question. Our finding that *C. elegans nrx-1* regulates the secretion of neuropeptides, if conserved in humans, would represent a novel mechanism by which neurexins alter neuronal function and behavior that would help understand pathophysiology of neurodevelopmental and neuropsychiatric disorders. This finding could also implicate new pathways and identify new targets to treat the behavioral changes associated with such conditions.

Neuropeptides are slower-acting, long-range, and promiscuous signaling molecules compared with classical neurotransmitters and signal via multiple neuropeptide receptors (GPCRs) that can in turn bind multiple NPs. This signaling flexibility allows NPs to influence multiple neuronal populations and circuits simultaneously, potentially contributing to the broad spectrum of phenotypes observed among individuals with autism. Neuropeptides, including oxytocin, vasopressin, and endogenous opioids, modulate social and behavioral processes in humans and have been investigated in clinical trials targeting autism-associated symptoms (Sikich et al. 2021; Borie, Theofanopoulou, and Andari 2021), while neurotensin, VIP, and CGRP are emerging as potential mechanistic and therapeutic targets (Angelidou et al. 2010; Tsilioni et al. 2019; Nelson et al. 2001).

**Supplementary Table 1. Evidence for neuropeptide expression in *C. elegans* motor neurons.**

**Supplementary Table 2. Strain and plasmid information.**

## MATERIALS AND METHODS

### *C. elegans* strain maintenance

all strains were maintained on Nematode Growth Medium (NGM) plates seeded with OP50 *Escherichia coli* at 20+C under standard conditions (Brenner 1974). All strains and mutant alleles included are listed in **Supplemental Table 2**. All experiments were performed on bleach synchronized day 1 adults unless otherwise noted.

### Cloning and constructs

All plasmids are listed in the **Supplementary Table 2** along with primer sequences for each promoter and the inserts for plasmids we generated. All plasmids were made by subcloning promoters, or cDNA inserts into plasmids by Epoch Life Science Inc., as described below. The *nrx-1(α)* and *nrx-1(*γ) transgene plasmids were generated by subcloning each promoter (*unc-129*p*, unc-47*p*, myo-3*p) to replace the *ric-19* promoter in pMPH34 (*ric-19*p*::sfGFP::nrx-1(α)*) or pMPH35 (*ric-19*p*::sfGFP::nrx-1(gamma)*). The *nlp-21*(ß-isoform) transgene plasmid (pMPHVT3) was generated by subcloning the *nlp-21* ß-isoform cDNA into pJQ355 (kind gift of Derek Sieburth), which already had the *unc-129* promoter and Venus tag. The *flp-9* transgene plasmids were generated by subcloning the *flp-9* cDNA into pMPHVT3 described above to replace the *nlp-21*(ß-isoform) and then subcloning the *unc-47* promoter to replace the *unc-129* promoter. The *ida-1* transgene plasmids were generated by first subcloning mRuby to replace the GFP in (pmdb466 (*srh-220p::ida-1::gfp*), a kind gift from Mario DeBono (Laurent et al. 2018)) then subcloning each promoter (*unc-129*p*, unc-47*p) to replace the *srh-220*p promoter (*unc-129*p*::ida-1::mRuby* and *unc-47*p*::ida-1::mRuby*).

### Generation of transgenes

*C. elegans* transformants were obtained as described previously (Mello and Fire 1995). A 100 ng/µl concentration injection mix was prepared with 50 ng of the transgene plasmid DNA, 5-15 ng/µl of co-injection marker (*myo-3*p*::RFP* or *unc-122*p*::GFP*) and carrier DNA (the 100 base pair gene ruler (NEB), at least 2-3 independent transgenic lines were characterized and analyzed to confirm expression levels and transmittance, after which a single line was selected for comprehensive analysis based on consistent expression levels and moderate to high transmittance.

### Microscopy

#### Sample preparation for confocal and Airyscan image acquisition

For microscopy, fresh 5% agarose pads were prepared. 10-30 day 1 bleach synchronized adult *C. elegans* were picked and placed onto agarose pads and the worms were let to be anesthetized using 4-5 µl of 100µM paralytic sodium azide for 5-10 minutes. After the worms stopped moving, a #1 thickness coverslip was placed on top of the *C. elegans.* After the slide preparation, they were then analyzed using a 63X objective on a Leica SP8 point scanning Confocal Microscope, with z-stack images taken at 0.6 μm spanning expression.

Confocal (non-Airyscan) images were acquired using a Leica TCS SP8 LAS X point-scanning confocal microscope equipped with a 63×/1.4 NA PL APO oil immersion objective. GFP was excited with a 488 nm laser at 1–3% laser power, and fluorescence emission was detected using a HyD2 detector over a wavelength range of 494–550 nm with the detector gain set to 40%. RFP was excited using a 552 nm laser at 0.6% laser power, and emission was collected using a HyD4 detector over a wavelength range of 552–784 nm with the detector gain set to 18.6%.GFP and RFP channels were acquired sequentially on a line-by-line basis using a pixel dwell time of 102.38 µs, a pinhole size of 1 Airy units (AU), 1-line averaging. The z-step size of 0.6 µm was taken for all images. Images were acquired at a format of 512 × 512 pixels using a scan speed of 700 Hz in bidirectional scanning mode with a zoom factor of 2.

Airyscan images were acquired on a Zeiss LSM 880 confocal (ZEN Black v. 2.3) managed by the CDB Microscopy Core at the University of Pennsylvania using a 63x/1.4 NA Plan Apochromat oil immersion objective. A 32-channel GaAsP Airyscan detector was used to collect fluorescence for both GFP and mCherry channels with an effective pinhole of 1.26 AU. For the GFP channel, excitation was provided by the 488 nm line of an argon-ion laser. Emission was selected using a combination of a 615 nm short-pass beamsplitter and double bandpass filter BP 420-480 + BP 495-550. For the mCherry channel, excitation was provided by a 561 nm DPSS laser and emission was selected using a combination of a 570 nm longpass beamsplitter and double bandpass filter BP420-480 + LP605. Channels were captured sequentially by frame with a pixel dwell time of 1.09 µs and no averaging. Pixel size was 0.043 µm. 5-slice z-stacks were acquired for all images using a z-step size of 0.185 µm (total z-stack range was 0.74 microns). After acquisition, ZEN Black was used for Airyscan processing of the raw 12-bit data; for all images, the filter strength was automatically set by the Zeiss algorithm.

#### Coelomocyte uptake assay

Coelomocyte uptake assays were performed as described previously (Sieburth, Madison and Kaplan, 2007; Hao et al., 2012). Z-stack images of the posterior coelomocytes were acquired from bleach-synchronized day 1 adult *C. elegans*. A region of interest (ROI) encompassing the entire coelomocyte was defined, and Z-stacks were converted into maximum-intensity projections (MIPs). Fluorescence intensity was quantified from the MIPs using ImageJ (NIH), normalized, and plotted using GraphPad Prism 11.

#### Synaptic markers

Line scans were acquired from worms positioned with the dorsal side of animals facing the objective. A region of interest (ROI) was defined, and fluorescence parameters, including mean fluorescence intensity, integrated density, puncta number, and colocalization index, were quantified for the dorsal nerve cord using ImageJ (NIH). For quantification of the UNC-31 reporter in the head region, an ROI encompassing the nerve ring was defined, and the total fluorescent area was measured following image thresholding in ImageJ.

#### Auxin (IAA)-inducible NRX-1 degradation

Tissue-specific knockdown of NRX-1 was achieved using the auxin-inducible degradation strategy as described previously (Sharma et al. 2024).Briefly, we crossed strains expressing auxin receptor, TIR1 (TRANSPORT INHIBITOR RESPONSE 1) either under neuronal (*rgef-1*) or muscle (*myo-3*) promoters and crossed with another strain expressing NRX-1 tagged with AID (auxin-inducible degron). Auxin Indole-3-acetic acid (IAA; ChemProducts) was added in NGM media during the pouring process. We tested at final concentrations of 1, 2 and 4 mM to evaluate bacterial lawn viability, worm growth and knockdown efficiency. A concentration of 1 mM IAA was selected for subsequent coelomocyte uptake assays, as it provided optimal worm growth while maintaining effective protein depletion. Degradation of the NRX-1::sfGFP::AID fusion protein could not be directly assessed because the overlapping NLP-21::Venus fluorescence signal precluded reliable quantification of the sfGFP signal, but previous characterization demonstrated strong loss of signal using this protocol (Bastien, Cowen, and Hart 2023).

#### Statistics

All statistical analyses and data visualization were performed using GraphPad Prism 11 (GraphPad Software). For the coelomocyte uptake assay, the fluorescence intensity of individual coelomocyte from each worm was quantified and plotted as mean ± SEM. Comparisons between two groups were performed using either an unpaired two-tailed Student’s *t*-test or Welch’s t-test, whereas comparisons among multiple groups were performed using one-way ANOVA followed by Tukey’s multiple-comparison test. Imaging was performed on at least three independent experimental days, with identical microscope acquisition settings maintained across all imaging sessions.

## Acknowledgments

The authors thank the Penn Worm Group labs for their feedback on this project, Hart lab members for their support and comments on the manuscript, and the CDB Microscopy Core (RRID SCR_022373) and Andrea Stout for help with Airyscan imaging. Some strains were provided by the CGC, which is funded by NIH Office of Research Infrastructure Programs (P40 OD010440).

## Data and Resource Availability

All relevant data and details of resources can be found within the article and its supplementary information and are available upon request.

## Author Contributions

VT and MPH conceived and designed the study and experiments, and VT conducted all experiments, processed, analyzed, and interpreted all data with help from MPH. VT wrote the manuscript with assistance from MPH.

## REFERENCES

Al Shehhi, Maryam, Eva B. Forman, Jacqueline E. Fitzgerald, et al. 2019. “NRXN1 Deletion Syndrome; Phenotypic and Penetrance Data from 34 Families.” European Journal of Medical Genetics 62 (3): 204–9.

AlHousni, Samira Said, Ahmed Idris, Watfa Al-Mamari, and Abeer Al-Sayegh. 2026. “Clinical Phenotypes Associated with NRXN1 Deletions in Five Children with Autism Spectrum Disorder in Oman.” Sultan Qaboos University Medical Journal 26 (1): 448–54.

Angelidou, Asimenia, Konstantinos Francis, Magdalini Vasiadi, et al. 2010. “Neurotensin Is Increased in Serum of Young Children with Autistic Disorder.” Journal of Neuroinflammation 7 (1): 48.

Ashley, Guinevere E., Tam Duong, Max T. Levenson, et al. 2021. “An Expanded Auxin-Inducible Degron Toolkit for Caenorhabditis Elegans.” Genetics 217 (3): iyab006.

Babu, Kavita, Zhitao Hu, Shih-Chieh Chien, Gian Garriga, and Joshua M. Kaplan. 2011. “The Immunoglobulin Super Family Protein RIG-3 Prevents Synaptic Potentiation and Regulates Wnt Signaling.” Neuron 71 (1): 103–16.

Bastien, Brandon L., Mara H. Cowen, and Michael P. Hart. 2023. “Distinct Neurexin Isoforms Cooperate to Initiate and Maintain Foraging Activity.” Translational Psychiatry 13 (1): 367.

Bono, M. de, and C. I. Bargmann. 1998. “Natural Variation in a Neuropeptide Y Receptor Homolog Modifies Social Behavior and Food Response in C. Elegans.” Cell 94 (5): 679–89.

Borie, Amelie M., Constantina Theofanopoulou, and Elissar Andari. 2021. “The Promiscuity of the Oxytocin-Vasopressin Systems and Their Involvement in Autism Spectrum Disorder.” Handbook of Clinical Neurology, Handbook of clinical neurology, vol. 182: 121–40.

Boucard, Antony A., Jaewon Ko, and Thomas C. Südhof. 2012. “High Affinity Neurexin Binding to Cell Adhesion G-Protein-Coupled Receptor CIRL1/Latrophilin-1 Produces an Intercellular Adhesion Complex.” The Journal of Biological Chemistry 287 (12): 9399–413.

Brenner, S. 1974. “THE GENETICS OF *CAENORHABDITIS ELEGANS*.” Genetics 77 (1): 71–94.

Cameli, Cinzia, Marta Viggiano, Magali J. Rochat, et al. 2021. “An Increased Burden of Rare Exonic Variants in NRXN1 Microdeletion Carriers Is Likely to Enhance the Penetrance for Autism Spectrum Disorder.” Journal of Cellular and Molecular Medicine 25 (5): 2459–70.

Cao, Xueshan, and Katsuhiko Tabuchi. 2017. “Functions of Synapse Adhesion Molecules Neurexin/Neuroligins and Neurodevelopmental Disorders.” Neuroscience Research 116 (March): 3–9.

Cheung, Amy, Kotaro Konno, Yuka Imamura, et al. 2023. “Neurexins in Serotonergic Neurons Regulate Neuronal Survival, Serotonin Transmission, and Complex Mouse Behaviors.” eLife 12 (e85058): e85058.

Cowen, Mara H., Dustin Haskell, Kristi Zoga, Kirthi C. Reddy, Sreekanth H. Chalasani, and Michael P. Hart. 2024. “Conserved Autism-Associated Genes Tune Social Feeding Behavior in C. Elegans.” Nature Communications 15 (1): 9301.

Engisch, K. L., and M. C. Nowycky. 1996. “Calcium Dependence of Large Dense-Cored Vesicle Exocytosis Evoked by Calcium Influx in Bovine Adrenal Chromaffin Cells.” The Journal of Neuroscience: The Official Journal of the Society for Neuroscience 16 (4): 1359–69.

Ferdos, Shima, Johannes Brockhaus, Markus Missler, and Astrid Rohlmann. 2021. “Deletion of β-Neurexins in Mice Alters the Distribution of Dense-Core Vesicles in Presynapses of Hippocampal and Cerebellar Neurons.” Frontiers in Neuroanatomy 15: 757017.

Fernando, Michael B., Yu Fan, Yanchun Zhang, et al. 2025. “Phenotypic Complexities of Rare Heterozygous Neurexin-1 Deletions.” Nature 642 (8068): 710–20.

Friedman, Amanda E., Michele Perni, Josephine Millard, Damjan Karanfilovski, Michael Granato, and Philip D. Campbell. 2026. “Unique and Overlapping Behavioral Effects of Isoform-Specific NRXN1 Deletions.” Disease Models & Mechanisms 19 (7). 10.1242/dmm.052862.

Fuccillo, Marc V., and Changhui Pak. 2021. “Copy Number Variants in Neurexin Genes: Phenotypes and Mechanisms.” Current Opinion in Genetics & Development 68 (June): 64–70.

Gomez, Andrea M., Lisa Traunmüller, and Peter Scheiffele. 2021. “Neurexins: Molecular Codes for Shaping Neuronal Synapses.” Nature Reviews. Neuroscience 22 (3): 137–51.

Gottmann, Kurt. 2008. “Transsynaptic Modulation of the Synaptic Vesicle Cycle by Cell-Adhesion Molecules.” Journal of Neuroscience Research 86 (2): 223–32.

Gracheva, Elena O., Anna O. Burdina, Denis Touroutine, Martine Berthelot-Grosjean, Hetal Parekh, and Janet E. Richmond. 2007. “Tomosyn Negatively Regulates CAPS-Dependent Peptide Release at Caenorhabditis Elegans Synapses.” The Journal of Neuroscience: The Official Journal of the Society for Neuroscience 27 (38): 10176–84.

Hammarlund, Marc, Oliver Hobert, David M. Miller 3rd, and Nenad Sestan. 2018. “The CeNGEN Project: The Complete Gene Expression Map of an Entire Nervous System.” Neuron 99 (3): 430–33.

Haskell, Dustin, William R. Haury, Myra Granato, Brandon L. Bastien, and Michael P. Hart. 2025. “Insertion of Rare Autism Variants in Synaptic Genes Induce Novel Behavioral Phenotypes in C. Elegans.” In bioRxivorg. bioRxiv, December 22. 10.64898/2025.12.19.695228.

Hu, Zhitao, Sabrina Hom, Tambudzai Kudze, et al. 2012. “Neurexin and Neuroligin Mediate Retrograde Synaptic Inhibition in C. Elegans.” Science (New York, N.Y.) 337 (6097): 980–84.

Khoja, Sheraz, Mulatwa T. Haile, and Lulu Y. Chen. 2023. “Advances in Neurexin Studies and the Emerging Role of Neurexin-2 in Autism Spectrum Disorder.” Frontiers in Molecular Neuroscience 16 (February): 1125087.

Kreutzberger, Alex J. B., Volker Kiessling, Binyong Liang, et al. 2017. “Reconstitution of Calcium-Mediated Exocytosis of Dense-Core Vesicles.” Science Advances 3 (7): e1603208.

Kurshan, Peri T., Sean A. Merrill, Yongming Dong, et al. 2018. “Γ-Neurexin and Frizzled Mediate Parallel Synapse Assembly Pathways Antagonized by Receptor Endocytosis.” Neuron 100 (1): 150–166.e4.

Lang, J., Y. Ushkaryov, A. Grasso, and C. B. Wollheim. 1998. “Ca2+-Independent Insulin Exocytosis Induced by Alpha-Latrotoxin Requires Latrophilin, a G Protein-Coupled Receptor.” The EMBO Journal 17 (3): 648–57.

Laurent, Patrick, Queelim Ch’ng, Maëlle Jospin, Changchun Chen, Ramiro Lorenzo, and Mario de Bono. 2018. “Genetic Dissection of Neuropeptide Cell Biology at High and Low Activity in a Defined Sensory Neuron.” Proceedings of the National Academy of Sciences of the United States of America 115 (29): E6890–99.

Li, Chris, and Kyuhyung Kim. 2010. “Neuropeptide Gene Families in Caenorhabditis Elegans.” Advances in Experimental Medicine and Biology (Boston, MA), Advances in experimental medicine and biology, vol. 692: 98–137.

Liu, Jiaxiang, Yanxuan Zhang, Ruijia Jin, Yuxin Zhu, and Jianrui Chen. 2026. “Roles of NRXN1 in Neuropsychiatric Disorders: From Genetic Lesion to Molecular Mechanism.” Frontiers in Neuroscience 20 (1808921): 1808921.

Luo, Fujun, Alessandra Sclip, Man Jiang, and Thomas C. Südhof. 2020. “Neurexins Cluster Ca2+ Channels within the Presynaptic Active Zone.” The EMBO Journal 39 (7): e103208.

Maro, Géraldine S., Shangbang Gao, Agnieszka M. Olechwier, et al. 2015. “MADD-4/Punctin and Neurexin Organize C. Elegans GABAergic Postsynapses through Neuroligin.” Neuron 86 (6): 1420–32.

Mello, C., and A. Fire. 1995. “DNA Transformation.” Methods in Cell Biology 48: 451–82.

Missler, Markus. 2003. “Synaptic Cell Adhesion Goes Functional.” Trends in Neurosciences 26 (4): 176–78.

Nelson, Karin B., Judith K. Grether, Lisa A. Croen, et al. 2001. “Neuropeptides and Neurotrophins in Neonatal Blood of Children with Autism or Mental Retardation.” Annals of Neurology 49 (5): 597–606.

Padmanabhan, Nirmala, Rebecca E. Twilley, Shinichiro Oku, et al. 2026. “Neurexin-1γ Drives Synaptic Transmission through Structural Integration of Neurotransmitter Release Machinery and Postsynaptic Receptor Nanodomains.” Neuron, ahead of print, July 21. 10.1016/j.neuron.2026.06.018.

Pandey, Pratima, Ashwani Bhardwaj, and Kavita Babu. 2017. “Regulation of WNT Signaling at the Neuromuscular Junction by the Immunoglobulin Superfamily Protein RIG-3 in Caenorhabditis Elegans.” Genetics 206 (3): 1521–34.

Philbrook, Alison, Shankar Ramachandran, Christopher M. Lambert, et al. 2018. “Neurexin Directs Partner-Specific Synaptic Connectivity in C. Elegans.” eLife 7 (July): e35692.

Röthe, Juliane, Doreen Thor, Jana Winkler, et al. 2019. “Involvement of the Adhesion GPCRs Latrophilins in the Regulation of Insulin Release.” Cell Reports 26 (6): 1573–1584.e5.

Rowen, Lee, Janet Young, Brian Birditt, et al. 2002. “Analysis of the Human Neurexin Genes: Alternative Splicing and the Generation of Protein Diversity.” Genomics 79 (4): 587–97.

Sasidharan, Nikhil, Marija Sumakovic, Mandy Hannemann, et al. 2012. “RAB-5 and RAB-10 Cooperate to Regulate Neuropeptide Release in Caenorhabditis Elegans.” Proceedings of the National Academy of Sciences of the United States of America 109 (46): 18944–49.

Selen, Ayşegül Tuğba Hıra, Necati Uzun, İbrahim Kılınç, and Ahmet Osman Kılıç. 2026. “Altered NCAM1, NRXN1, NLGN4, and CDH2 Levels in Individuals with Autism Spectrum Disorder and Their Unaffected Siblings.” Clinical Psychopharmacology and Neuroscience: The Official Scientific Journal of the Korean College of Neuropsychopharmacology, ahead of print, June 24. 10.9758/cpn.26.1398.

Shahi, Navneet, Shruti Thapliyal, and Kavita Babu. 2025. “Sensory Modulation of Neuropeptide Signaling by CASY-1 Gates Cholinergic Transmission at Caenorhabditis Elegans Neuromuscular Junction.” Journal of Biosciences 50: 4.

Sharma, Nidhi, Filipe Marques, and Paschalis Kratsios. 2024. “Protocol for Auxin-Inducible Protein Degradation in C. Elegans Using Different Auxins and TIR1-Expressing Strains.” STAR Protocols 5 (3): 103133.

Sieburth, Derek, Jon M. Madison, and Joshua M. Kaplan. 2007. “PKC-1 Regulates Secretion of Neuropeptides.” Nature Neuroscience 10 (1): 49–57.

Sikich, Linmarie, Alexander Kolevzon, Bryan H. King, et al. 2021. “Intranasal Oxytocin in Children and Adolescents with Autism Spectrum Disorder.” The New England Journal of Medicine 385 (16): 1462–73.

Suckow, Arthur T., Davide Comoletti, Megan A. Waldrop, et al. 2008. “Expression of Neurexin, Neuroligin, and Their Cytoplasmic Binding Partners in the Pancreatic Beta-Cells and the Involvement of Neuroligin in Insulin Secretion.” Endocrinology 149 (12): 6006–17.

Tikiyani, Vina, Lei Li, Pallavi Sharma, Haowen Liu, Zhitao Hu, and Kavita Babu. 2018. “Wnt Secretion Is Regulated by the Tetraspan Protein HIC-1 through Its Interaction with Neurabin/NAB-1.” Cell Reports 25 (7): 1856–1871.e6.

Tromp, Alisha, Bryan Mowry, and Jean Giacomotto. 2021. “Neurexins in Autism and Schizophrenia-a Review of Patient Mutations, Mouse Models and Potential Future Directions.” Molecular Psychiatry 26 (3): 747–60.

Tsilioni, Irene, Arti B. Patel, Harry Pantazopoulos, et al. 2019. “IL-37 Is Increased in Brains of Children with Autism Spectrum Disorder and Inhibits Human Microglia Stimulated by Neurotensin.” Proceedings of the National Academy of Sciences of the United States of America 116 (43): 21659–65.

Xia, Xiaofeng, Volkmar Lessmann, and Thomas F. J. Martin. 2009. “Imaging of Evoked Dense-Core-Vesicle Exocytosis in Hippocampal Neurons Reveals Long Latencies and Kiss-and-Run Fusion Events.” Journal of Cell Science 122 (Pt 1): 75–82.

Zahn, Tobias R., Joseph K. Angleson, Margaret A. MacMorris, et al. 2004. “Dense Core Vesicle Dynamics in Caenorhabditis Elegans Neurons and the Role of Kinesin UNC-104.” Traffic (Copenhagen, Denmark) 5 (7): 544–59.

Zhong, Yuanxin, Li An, Yufeng Wang, Li Yang, and Qingjiu Cao. 2022. “Functional Abnormality in the Sensorimotor System Attributed to NRXN1 Variants in Boys with Attention Deficit Hyperactivity Disorder.” Brain Imaging and Behavior 16 (3): 967–76.

